# Lack of p62 impairs glycogen aggregation and exacerbates pathology in a mouse model of myoclonic epilepsy of Lafora

**DOI:** 10.1101/2021.06.03.446965

**Authors:** Pasquale Pellegrini, Arnau Hervera, Olga Varea, M. Kathryn Brewer, Iliana López-Soldado, Anna Guitart, Mònica Aguilera, Neus Prat, José Antonio del Río, Joan J. Guinovart, Jordi Duran

## Abstract

**Background:** Lafora disease (LD) is a fatal childhood-onset dementia characterized by the extensive accumulation of glycogen aggregates—the so-called Lafora Bodies (LBs)—in several organs. The accumulation of LBs in the brain underlies the neurological phenotype of the disease. LBs are composed of abnormal glycogen and various associated proteins, including p62, an autophagy adaptor that participates in the aggregation and clearance of misfolded proteins.

**Methods:** To study the role of p62 in the formation of LBs and its participation in the pathology of LD, we generated a mouse model of the disease (malin^KO^) lacking p62.

**Results:** Deletion of p62 prevented LB accumulation in skeletal muscle and cardiac tissue. In the brain, the absence of p62 altered LB morphology and increased susceptibility to epilepsy.

**Conclusions:** These results demonstrate that p62 participates in the formation of LBs and suggest that the sequestration of abnormal glycogen into LBs is a protective mechanism through which to reduce the deleterious consequences of its accumulation in the brain.

## BACKGROUND

Glycogen, a branched polymer of glucose, is found in most tissues but is particularly abundant in the liver and muscle (1). In the brain, glycogen is present mainly in astrocytes (2–4), although neurons also have an active glycogen metabolism that contributes to their function (5, 6). In mammals, glycogen is synthesized by glycogen synthase and degraded by glycogen phosphorylase. The muscular isoform of glycogen synthase (MGS) is expressed in most tissues, including the brain.

Progressive myoclonic epilepsy type 2 (EPM2, OMIM #254780) or Lafora disease (LD) is an autosomal recessive disease characterized by severe and progressive myoclonus epilepsy, and neurodegeneration rapidly progressing to dementia and death within 5-10 years after the onset (7, 8). LD is caused by mutations in either the *EPM2A* gene, which encodes laforin, a dual specificity phosphatase with a carbohydrate-binding domain, or *EPM2B* (also *NHLRC1*), which encodes malin, an E3-ubiquitin ligase. The histopathological and clinical outcomes of LD patients and mouse models of LD carrying mutations in either of these two genes are very similar, thereby indicating that malin and laforin participate in the same physiological process. The hallmark of the disease is the accumulation of cytoplasmic aggregates of poorly branched glycogen called Lafora bodies (LBs) in several tissues (9, 10). In the brain, LBs are found in astrocytes and neurons (11–13). Neuronal LBs (nLBs) typically manifest as single, large, round and juxtanuclear aggregates, while astrocytic LBs are smaller and amorphous and have a granular distribution throughout astrocytic processes (12). We refer to these astrocytic LBs as CAL *(corpora amylacea-like*), since they are morphologically similar to *corpora amylacea—*glycogen aggregates that accumulate in normal aging (8, 14). Blocking or reducing brain glycogen synthesis in LD mouse models prevents the progression of the disease (13, 15–18), thereby indicating that glycogen accumulation underlies the pathophysiology of LD. Furthermore, forced accumulation of glycogen in neurons leads to neuronal loss (19) while in astrocytes it induces neuroinflammation (13).

In addition to glycogen, LBs contain a number of proteins, including laforin (in malin-deficient LD), enzymes involved in glycogen metabolism such as MGS, ubiquitinated proteins, and the autophagy adaptor p62 (17, 18, 20). The presence of ubiquitin and p62 suggests that, like other insoluble molecular aggregates characteristic of neurodegenerative diseases, LBs could be targets for autophagic clearance (17, 20–23). In this regard, the mechanisms that drive the formation and clearance of LBs have not been identified yet. Since p62 has been shown to aggregate polyubiquitinated proteins (24), it could play a similar role in the formation of LBs. Furthermore, although glycogen accumulation underlies LD pathogenesis, it remains to be determined whether the sequestration of this polysaccharide into LBs is protective (to minimize the toxic consequences of the accumulation of abnormal glycogen) or pathogenic (LBs themselves being the toxic species). Finally, the accumulation of p62 *per se* is deleterious for neurons (25, 26) and other cell types (27, 28). In this regard, p62 depletion clears nuclear inclusion bodies and increases lifespan in a model of Huntington’s disease (29).

Given all of the above, p62 may exert a neuroprotective or a neurotoxic function in the context of LD. To study the contribution of p62 to LB formation and to the pathophysiology of LD, we generated a malin knockout mouse (malin^KO^) (30) devoid of p62 (malin^KO^+p62^KO^). Our results demonstrate that p62 is essential for LB formation in skeletal muscle and cardiac tissue. In the brain, p62 is also involved in the formation of these aggregates. When this protein is absent in this organ, neuroinflammation is mildly enhanced and susceptibility to epilepsy is exacerbated. These observations identify p62 as a key player in the cellular protective response against glycogen aggregates.

## MATERIALS AND METHODS

### Animal studies

All procedures were approved by the Barcelona Science Park’s Animal Experimentation Committee and were carried out following Spanish (BOE 34/11370–421, 2013) and European Union (2010/63/EU) regulations, and The National Institutes of Health guidelines for the care and use of laboratory animals. For the generation of the malin^KO^ + p62^KO^ model, malin^KO^ mice (30) were crossed with p62^KO^ animals (23). After weaning at 3 weeks of age, tail clippings were taken for genotyping by qPCR (performed by TransnetYX). Experiments were conducted using littermates, and males and females were included in each group. Mice were maintained on a 12/12 h light/dark cycle under specific pathogen-free conditions in the Animal Research Center (Barcelona Science Park) and allowed free access to a standard chow diet and water.

### Glycogen quantification

Mice were deeply anesthetized and decapitated. Whole brains and quadriceps were quickly removed, frozen, and pulverized in liquid nitrogen. For glycogen measurements, frozen tissue aliquots were boiled in 30% KOH for 15 min and glycogen was precipitated in 60% ethanol and then determined by an amyloglucosidase-based assay (5).

### Western blot

For western blot, lysates of frozen tissue aliquots were prepared using the following buffer: 25 mM Tris-HCl (pH 7.4), 25 mM NaCl, 1% Triton X-100, 0.1% SDS, 0.5 mM EGTA, 10 mM sodium pyrophosphate, 1 mM sodium orthovanadate, 10 mM NaF, 25 nM okadaic acid and a protease inhibitor cocktail tablet (Roche). Soluble and insoluble fractions of total homogenates were obtained as previously described (30). Briefly, total homogenates were centrifuged at 13,000 rpm for 15 min at 4°C. The pellet containing the insoluble fraction was resuspended in the same volume as the supernatant corresponding to the soluble fraction. Samples were loaded on 10% acrylamide gels for SDS-PAGE and transferred to Immobilon membranes (Millipore). The following primary antibodies were used: anti-MGS (3886, Cell Signaling); anti-laforin (3.5.5, kindly provided by Dr. Santiago Rodríguez de Córdoba); and anti-p62 (GP62-C, Progen). The following secondary antibodies were used: anti-rabbit and anti-mouse IgG-HRP (GE Healthcare); and anti-guinea pig HRP (Jackson Immuno Research). Proteins were detected by the ECL method (Immobilon Western Chemiluminescent HRP Substrate, Millipore), and loading control of the western blot membrane was performed using the Revert total protein stain (LI-COR Bioscience).

### Histology and immunohistochemistry

Animals were deeply anesthetized and perfused transcardially with phosphate-buffered saline (PBS) containing 4% paraformaldehyde (PBS 4% PFA). Brains, skeletal muscles and hearts were removed, post-fixed overnight with PBS 4% PFA and embedded in paraffin blocks. Periodic acid-Schiff staining (PAS) was performed using an Artisanlink Pro machine (AR16511-2 kit, Dako-Agilent).

For immunohistochemistry, 3 μm paraffin-embedded tissue sections were either dewaxed and subjected to antigen retrieval treatment with Tris-EDTA buffer pH 9 for 20 min at 97°C using a PT Link (Dako – Agilent) or dewaxed as part of the antigen retrieval process using the Low pH EnVision ™ FLEX Target Retrieval Solutions (K8005, Dako-Agilent) for 20 min at 97°C using a PT Link (Dako – Agilent). Endogenous peroxidase was quenched with Peroxidase-Blocking Solution (S2023, Dako-Agilent). Non-specific binding was blocked using 5% of normal goat serum (16210064, Life technology) with 2.5% BSA (10735078001, Sigma) for 60 min. Also, unspecific endogenous mouse Ig staining was blocked using the Mouse on mouse (M.O.M) Immunodetection Kit (BMK-2202, Vector Laboratories). Primary mouse IgG1 anti-GFAP (MAB360, Merck Millipore) and rabbit pAb 1 anti-IBA1 (019-19741, WAKO) antibodies were diluted at 1:250 and 1:1000 respectively with EnVision FLEX Antibody Diluent (K800621, Dako-Agilent) and incubated overnight at 4°C. Tissue sections were then incubated for 45 min with Polyclonal Anti-Mouse 1:100 (P0447, Dako-Agilent) or a BrightVision Poly-HRP-Anti Rabbit IgG, RTU (Immunologic, DPVR-110HRP). Antigen–antibody complexes were revealed with 3-3’-diaminobenzidine (K3468, Dako). Sections were counterstained with haematoxylin (Dako, S202084) and mounted with Toluene-Free Mounting Medium (CS705, Dako) using a Dako CoverStainer.

For immunofluorescence, endogenous peroxidase was quenched by 10 min of incubation with Peroxidase-Blocking Solution (S2023, Dako-Agilent). Non-specific binding was blocked using 5% of normal goat serum (16210064, Life technology) with 2.5% BSA (10735078001, Sigma) for 60 min. Also, unspecific endogenous mouse Ig staining was blocked using the M.O.M Immunodetection Kit (BMK-2202, Vector Laboratories). Primary antibodies anti-MGS (15B1) (1:250, 3886 Cell Signaling), anti-GFAP (1:250, MAB360, Merck Millipore), anti-β-Tubulin III (1:500, T86660, Sigma Aldrich) and C3d antibody (1: 100, AF2655, R&D) were incubated overnight at 4°C.

The following secondary antibodies were used: an Alexa Fluor® 488 anti-mouse IgG (405319, BioLegend); Alexa Fluor® 488 anti-mouse IgG1 (A21121, ThermoFisher); Alexa Fluor 568® anti-mouse IgG2b (A21144, ThermoFisher); DyLight 594 anti-rabbit (DI1094, VectorLabs); or an Alexa Fluor 647® anti-rabbit IgG (A32733, ThermoFisher), diluted at 1:500 and incubated for 60 min. Samples were stained with DAPI (D9542, Sigma) and mounted with Fluorescence mounting medium (S3023 Dako). Specificity of staining was respectively confirmed by staining with rabbit IgG, polyclonal Isotype Control (ab27478, Abcam), mouse IgG1, Kappa Monoclonal (NCG01) Isotype Control (ab81032, Abcam) or a mouse IgM (PFR-03) Isotype Control (A1-10438, ThermoFisher).

Brightfield and fluorescent images were acquired with a NanoZoomer-2.0 HT C9600 digital scanner (Hamamatsu) equipped with a 20x objective. For super-resolution microscopy, images were acquired in a Zeiss 880 confocal microscope equipped with Fast Airyscan and a piezo-stage. A 63x magnification 1.40 NA oil-immersion lens with a digital zoom of 1.5x was used. The Z-step between the stacks was set at 0.8 μm. Fast Airyscan raw data were pre-processed with the automatic setting of Zen Black.

The stainings were analyzed by the digital software analysis package QuPath (31). For detection of the morphological features of nLBs and CAL, MGS-positive granules were identified using the Cell detection plugin (QuPath). Intensity thresholds were set for GFAP and βIII-Tubulin in the surrounding area of each MGS-positive granule. Morphology and intensity data were then exported and plotted with RStudio (32).

### RT-qPCR analyses

Total RNA of pulverized brains was prepared with the RNeasy Micro Kit (Qiagen), following the manufacturer’s instructions. Single-stranded complementary DNA was produced by reverse transcription using 1 μg of DNA-free RNA in a 20-μL reaction qScript cDNA SuperMix (Quanta bio). Quantitative polymerase chain reaction (PCR) was performed using SYBR green (Quanta bio) on the QuantStudio 6 Flex as per the manufacturer’s instructions. The ΔCt was defined as the difference between the Q-PCR cycles of the housekeeping gene and those of the target genes.

### Assessment of kainate-induced epilepsy

Mice were weighed and placed in individual cages to prevent contact between animals, which could startle them. They were then administered three consecutives intraperitoneal (i.p.) injections (6 mg/kg body weight) of the glutamate agonist kainic acid (KA) (Sigma) dissolved in 0.1 M PBS pH 7.4, in order to induce non-lethal convulsive seizures. Seizure intensity after KA injections was evaluated as described previously (33–35) for 240 minutes from the first KA administration. After the first KA injections, the animals developed hypoactivity and immobility (Grade I–II). After successive injections, hyperactivity (Grade III) and scratching with mild non-convulsive seizures (Grade IV) were often observed. Some animals progressed to a whole-body convulsive seizure with loss of balance control (Grade V). Extreme behavioral manifestations such as uncontrolled hopping activity or “popcorn behavior” (blinking seizure), as well as continuous or chronic seizures (> 1’ without body movement control) were included in Grade VI. All behavioral assessments were performed blind to the experimental group (genotype) in situ and were also recorded and reanalyzed blind to the first analysis.

### Statistical analysis

Two-group hypothesis testing was evaluated using an independent sample t-test performed with the GraphPad Prism software (La Jolla, CA, USA). Two-way analysis of variation (ANOVA) was used for comparing three or more groups. Data are represented as mean ± standard error of the mean (SEM). When indicated, linear mixed-effects model was used as follows.

#### nLB and CAL morphology

For each morphological parameter independently, linear mixed effect models were fitted with the R package lmerTest (36) using the parameters response variable, cell type, and the interaction between cell type and genotype as covariates of interest, the experimental group and the position as adjusting factors, and both the Mouse ID and the interaction between the Mouse ID and cell type as random effect to account for non-independence among data from the same mice. Transformations: for parameter values between 0 and 1, a logit transformation was applied. Otherwise, for parameter values larger than zero, a log transformation was considered.

#### Q-PCR analysis at gene level

For every gene, independently, linear mixed-effects models were fitted with the R package lmerTest using the ΔCt as response variable, Genotype as covariate of interest, experimental group as adjusting factor and Mouse ID as random effect to account for the variability of technical replicates. Adjustment for multiple testing (single-step correction method) was performed using the R package multcomp (37).

#### Q-PCR analysis for pro- and anti-inflammatory differences

For each mouse, ΔCt levels for the measured replicates were averaged out. For each gene, the mouse group (batch) effect was also balanced out and, finally, standardization was applied to these balanced data. The average value of each genotyping condition is shown in the graphs.

To evaluate consistency between gene patterns across inflammatory groups, a linear mixed-effects model was fitted with the R package lmerTest (36) using the interaction between the genotype and the inflammatory group as fixed effects and the gene Id as random effect. Adjustment for multiple testing (single-step correction method) was performed using the R package multcomp (37).

## RESULTS

### p62 progressively accumulates with age in the brain and skeletal muscle of malin^KO^ mice

We previously described increased levels of p62 in 11-month-old malin^KO^ brains (17, 18). The accumulated p62 is bound to LBs, both CAL and nLBs (12). To further characterize p62 accumulation in malin^KO^ tissues, we performed immunofluorescence against this protein. We observed an increase in p62 in the cortex and the hippocampus of 4- and 11-month-old malin^KO^ mice (Figure 1A), which paralleled the accumulation of LBs. Indeed, the number of p62-positive aggregates more than doubled in the brains of 11-month-old animals compared to those aged 4 months (Figure 1B). A massive increase in p62-positive aggregates was detected in the skeletal muscles of 11-month-old malin^KO^ animals compared to 4-month-old counterparts (Figure 1A, C).

**Figure 1.**
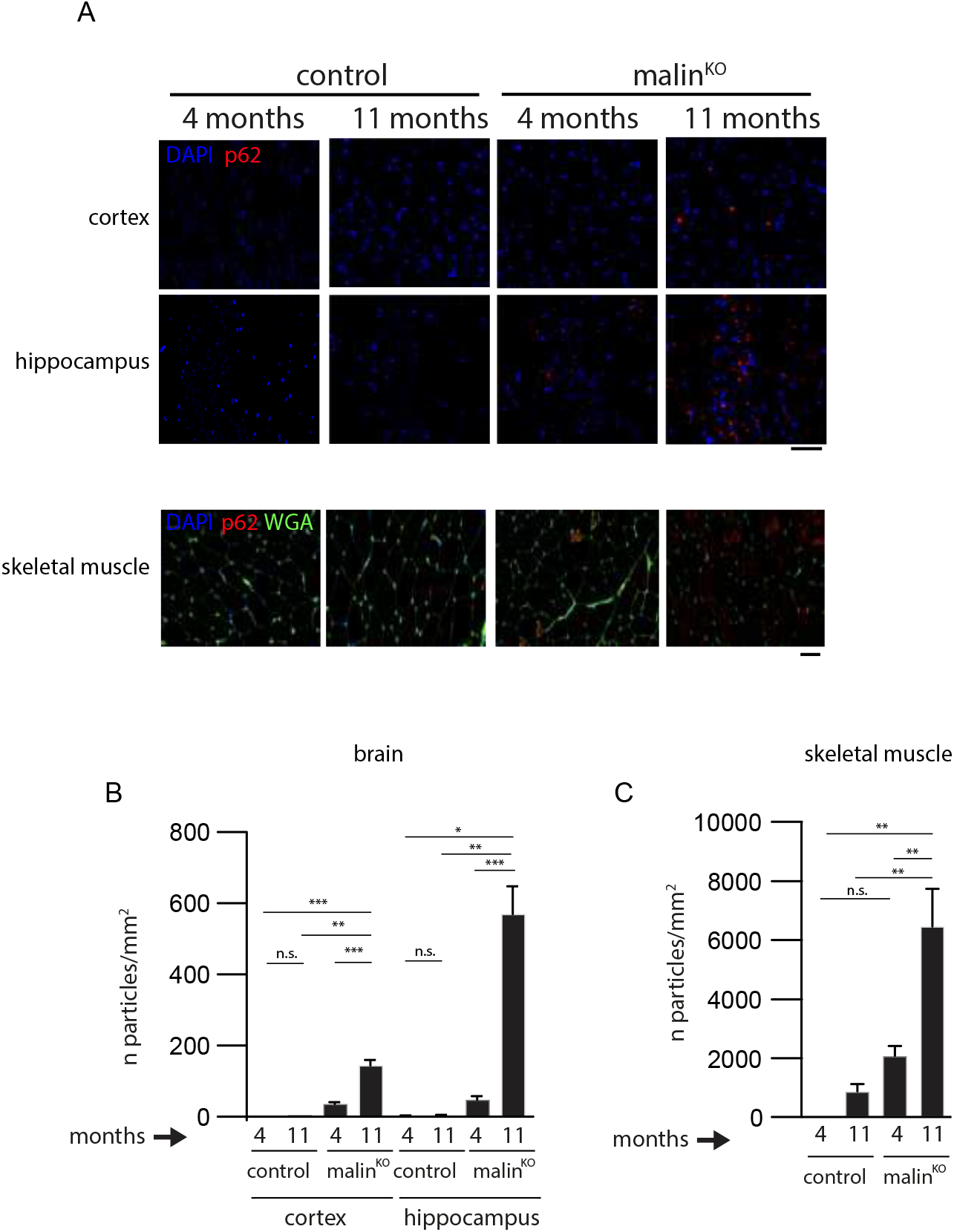
Accumulation of p62 aggregates over time in malin^KO^ mice. A. Representative images of the progressive accumulation of p62 in the cortex and hippocampus of malin^KO^ mice. Scale bar: 50 μm. B-C. Quantification of the number of p62+ LBs per area (n of particles/mm^2^) in the prefrontal cortex or hippocampus (B) and skeletal muscle (C). n=7-12 mice. For comparisons between groups, one-way ANOVA was performed using Prism7 software (GraphPad). * P<0.05, ** P<0.01, *** P<0.001, **** P<0.0001. *Adjusted p-values (cortex)*: control 4 m vs. control 11 m p=0.9997; control 4 m vs. malin^KO^ 4 m p=0.7191; control 4 m vs. malin^KO^ 11 m p=0.0010; control 11 m vs. malin^KO^ 4 m p=0.7829; control 11 m vs. malin^KO^ 11 m p=0.0013; and malin^KO^ 4 m vs. malin^KO^ 11 m p=0.0001. *Adjusted p-values (hippocampus)*: control 4 m vs. control 11 m p>0.9999; control 4 m vs. malin^KO^ 4 m p= 0.9929; control 4 m vs. malin^KO^ 11 m p=0.0109; control 11 m vs. malin^KO^ 4 m p=0.9936; control 11 m vs. malin^KO^ 11 m p=0.0111; and malin^KO^ 4 m vs. malin^KO^ 11 m p=0.0005. *Adjusted p-values (skeletal muscle)*: control 4 m vs. control 11 m p=0.9556; control 4 m vs. malin^KO^ 4 m p=0.5429; control 4 m vs. malin^KO^ 11 m p=0.0020; control 11 m vs. malin^KO^ 4 m p=0.7784; control 11 m vs. malin^KO^ 11 m p=0.0014; and malin^KO^ 4 m vs. malin^KO^ 11 m p=0.0041.

### p62 is essential for LB formation in muscle and heart tissue but not in the brain

To evaluate the impact of p62 deletion on LB formation and LD progression, we generated malin^KO^ mice devoid of p62 (malin^KO^+p62^KO^). The presence of LBs was visualized by periodic acid-Schiff staining (PAS), which specifically stains carbohydrates, and by immunofluorescence using anti-MGS antibody, since MGS is attached to LBs and can thus be used as an LB marker (13, 18).

The 11-month-old mice malin^KO^ animals showed abundant PAS-positive LBs in all the tissues analyzed, i.e. skeletal muscle, heart and brain, as previously described (17, 18, 30) (Figure 2A, Supplemental Figure 1A). Strikingly, the skeletal muscles of malin^KO^+p62^KO^ mice were devoid of PAS-positive aggregates (Figure 2A) and instead showed a diffused pattern of PAS-positive material. MGS immunostaining confirmed the absence of glycogen aggregates in the skeletal muscle of malin^KO^+p62^KO^ mice (Figure 2B). Similar results were observed in cardiac tissue, where deletion of p62 resulted in the absence of PAS-positive and MGS-positive aggregates (Supplemental Figure 1A, B). Although devoid of LBs, malin^KO^+p62^KO^ skeletal muscles showed an increase in total glycogen, as determined by biochemical quantification, similar to that seen in malin^KO^ muscles (Figure 2D). These results indicate that the skeletal muscles of malin^KO^+p62^KO^ mice also accumulated glycogen, although not in the form of LBs.

**Figure 2.**
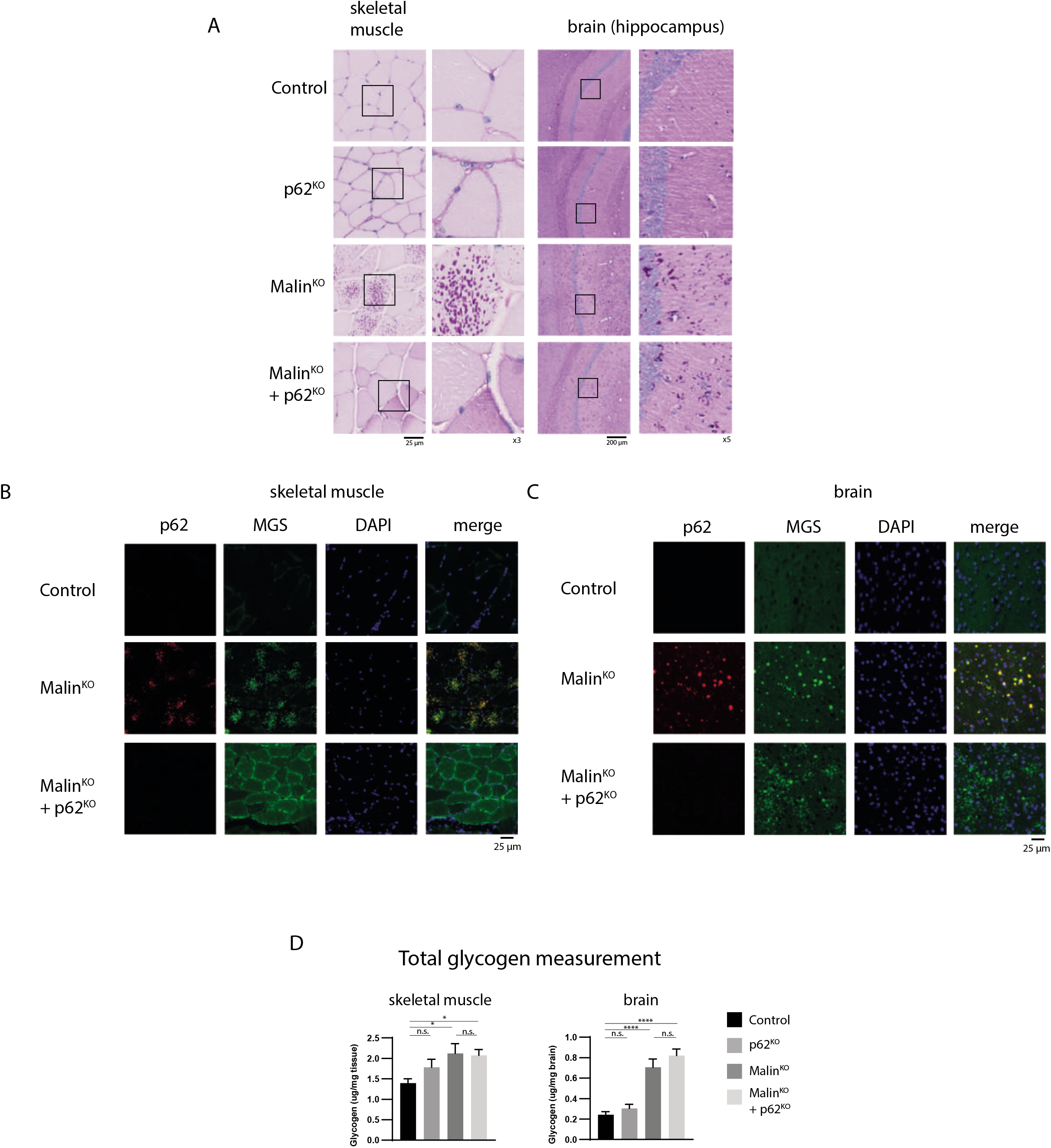
p62 deletion prevents glycogen aggregation in skeletal muscle but not in brain. A. Histological localization of LBs. Periodic acid-Schiff staining (PAS) is shown for the indicated tissues of 11-month-old malin^KO^ and malin^KO^+p62^KO^ mice. Scale bar= 25 μm (skeletal muscle and heart), 200 μm (brain). Brain aggregates are visible in the insets. B-C. Representative immunostaining of p62 and MGS in skeletal muscle (quadriceps) and brain (cortex). Scale bar=25 μm D. Glycogen content. For comparisons between groups, one-way ANOVA followed by Holm’s Multiple Comparisons Test was performed using Prism7 software (GraphPad). Results are presented as the group mean ± SEM. (n=6 animals per group). * P<0.05, ** P<0.01, *** P<0.001, **** P<0.0001. *Adjusted p-values.* Brain glycogen: control vs. p62KO p=0.4593; control vs. malin^KO^ p<0.0001; control vs. malin^KO^+p62^KO^ p<0.0001; and malin^KO^ vs. malin^KO^+p62^KO^ p= 0.3052. Muscle glycogen: control vs. p62^KO^ p= 0.3238; control vs. malin^KO^ p=0.0358; control vs. malin^KO^+p62^KO^ * p<0.05; and malin^KO^ vs. malin^KO^+p62^KO^ p=0.8546.

In contrast, the brains of these mice showed PAS-positive aggregates (Figure 2A). As expected, these aggregates contained MGS but not p62 (Figure 2C). Total glycogen levels were similarly elevated in malin^KO^+p62^KO^ and malin^KO^ brains (Figure 2D).

### Accumulation of insoluble MGS and laforin is prevented in the skeletal muscle of malin^KO^+p62^KO^ animals but not in the brain

Using western blot of total tissue homogenates, we next studied the content of proteins known to accumulate in malin^KO^ tissues. Since LBs are insoluble aggregates that precipitate under low-speed centrifugation (30), we also examined the distribution of these proteins between the soluble and insoluble fractions, the latter corresponding to the LB-enriched fraction. Western blots confirmed the absence of p62 in tissues from p62^KO^ and malin^KO^+p62^KO^ mice. In the skeletal muscles of malin^KO^ mice, MGS, laforin and p62 were significantly increased in the insoluble fraction (Figure 3A, B), as previously reported (18). However, in malin^KO^+p62^KO^ skeletal muscles, MGS and laforin showed levels similar in the insoluble fraction to those of controls (Figure 3A, B). These results were consistent with the absence of LBs in malin^KO^+p62^KO^ skeletal muscles.

**Figure 3.**
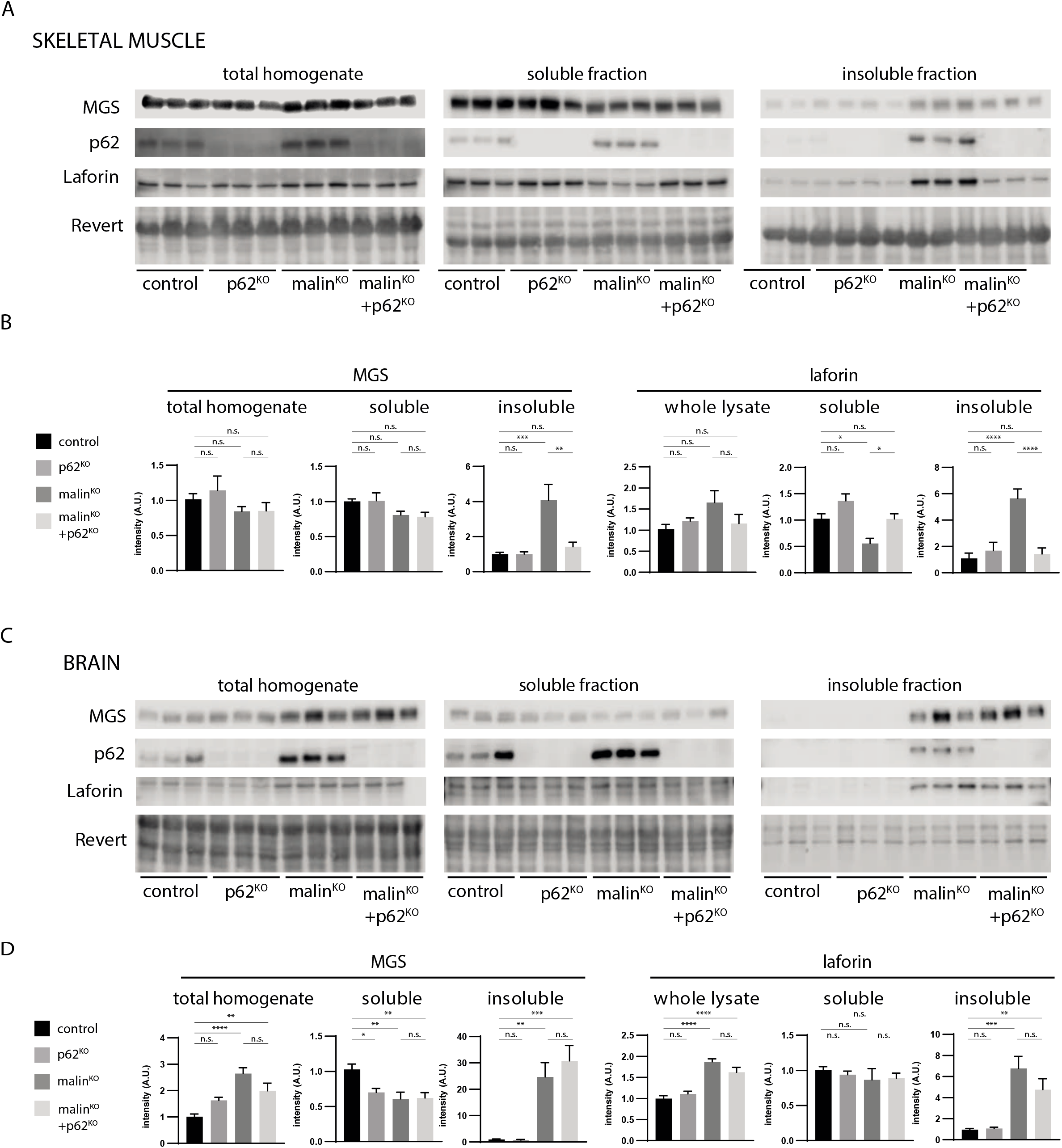
p62 deletion rescues accumulation of insoluble glycogen-bound proteins MGS and laforin in skeletal muscle but not in brain. A-C. Western blotting for MGS, p62 and laforin in muscle (A) and brain (C). Total protein was used as loading control. B-D. Densitometry of the western blots in skeletal muscle (B) and brain (D). For comparisons between groups, one-way ANOVA followed by Holm’s Multiple Comparisons Test was performed. N=6-9 animals/group were analyzed. Results are presented as the group mean ± SEM. * P<0.05, ** P<0.01, *** P<0.001, **** P<0.0001. *Adjusted p-values for MGS (a) or laforin (b)*: Brain whole lysates: control vs. p62^KO^ p=0.057 (a), p=0.3489 (b); control vs. malin^KO^ p<0.0001 (a) p<0.0001 (b); control vs. malin^KO^+p62^KO^ p=0.0066 (a), p<0.0001 (b); and malin^KO^ vs. malin^KO^+p62^KO^ p=0.057 (a), p=0.0898 (b). Brain soluble: control vs. p62^KO^ p=0.0143 (a), p=0.8557 (b); control vs. malin^KO^ p=0.0041 (a), p=0.7614 (b); control vs. malin^KO^+p62^KO^ p=0.0041 (a), p= 0.7614 (b); and malin^KO^ vs. malin^KO^+p62^KO^ p=0.9128 (a), p=0.8737 (b). Brain insoluble: control vs. p62^KO^ p=0.9649 (a), p=0.9179 (b); control vs. malin^KO^ p=0.0013 (a), p=0.0001 (b); control vs. malin^KO^+p62^KO^ p=0.0001 (a), p=0.0075 (b); and malin^KO^ vs. malin^KO^+p62^KO^ p=0.5051 (a), p=0.1591 (b). Muscle whole lysate: control vs. p62^KO^ p=0.8217 (a), p=0.7297 (b); control vs. malin^KO^ p=0.8217 (a), p=0.0919 (b); control vs. malin^KO^+p62^KO^ p=0.8217 (a). p=0.7297 (b); and malin^KO^ vs. malin^KO^+p62^KO^ p=0.9765 (a), p=0.1987 (b). Muscle soluble: control vs. p62^KO^ p=0.9597 (a), p=0.0573 (b); control vs. malin^KO^ p=0.1964 (a), p=0.0135 (b); control vs. malin^KO^+p62^KO^ p=0.1543 (a), p=0.9727 (b); and malin^KO^ vs. malin^KO^+p62^KO^ p=0.9597 (a), p=0.0135 (b). Muscle insoluble: control vs. p62^KO^ p=0.9966 (a), p=0.6862 (b); control vs. malin^KO^ p=0.0003 (a), p<0.0001 (b); control vs. malin^KO^+p62^KO^ p=0.7838 (a), p=0.6862 (b); and malin^KO^ vs. malin^KO^+p62^KO^ p=0.0013 (a) p<0.0001 (b).

In brain total homogenates, malin^KO^ mice showed an increase in MGS and laforin, which corresponded to an increase in the insoluble fraction (Figure 3C, D), as we previously described (18, 30). In contrast to skeletal muscles, the brains of malin^KO^+p62^KO^ mice showed similar increases in these two proteins in the insoluble fraction, consistent with the presence of LBs in this tissue.

### p62 depletion alters the morphology of brain LBs

Although LBs were still present in malin^KO^+p62^KO^ brains, we studied the morphology of these aggregates formed in the absence of p62. Super-resolution microscopy revealed that nLBs in malin^KO^+p62^KO^ animals appeared less dense and more irregular than the typical round, compact nLBs found in malin^KO^ brains (Figure 4A). The morphology of astrocytic LBs (CAL), which are inherently more heterogeneous than nLBs (12), was indistinguishable between the two genotypes, both showing highly irregular shapes (Figure 4B).

**Figure 4.**
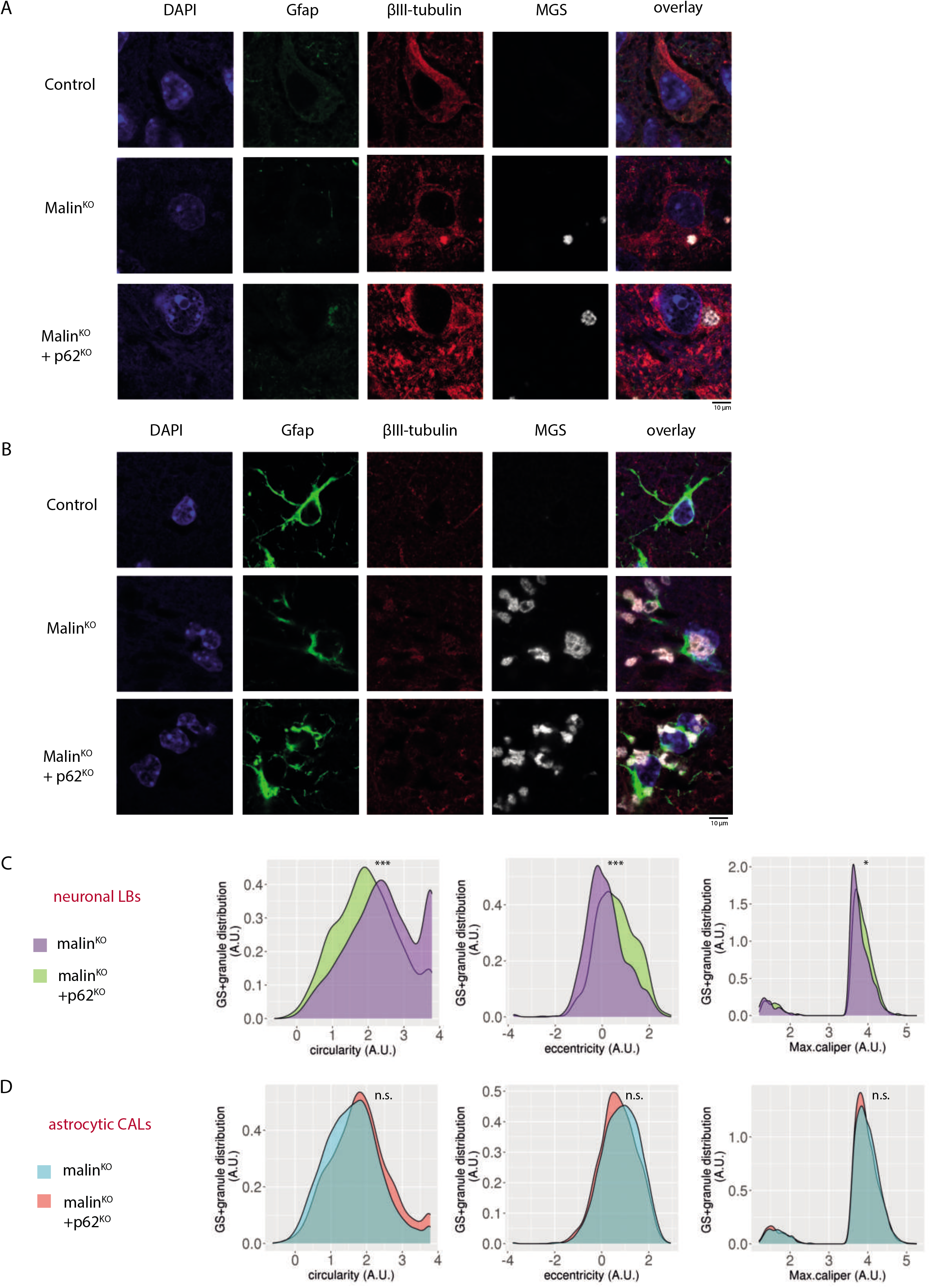
p62 deletion disrupts nLB but not CAL morphology. A-B. Representative examples of reconstructed super-resolution images of neuronal LBs (A) and astrocytic CAL (B) are shown. PFA-fixed tissue sections were incubated with anti-glycogen synthase (MGS), anti-GFAP and βIII-tubulin antibodies, nuclei were stained with Hoechst 33342. Scale bar: 10 μm. C-D. Quantifications of circularity (left panels), eccentricity (middle panels) and max caliper (right panels) in neuronal LBs (C) and astrocytic CAL (D). Linear mixed model analysis was performed for statistical analysis. N=6 animals per genotype. *Adjusted p-values* Circularity: astrocytic p=0.075; neuronal p<0.0001. Eccentricity: astrocytic p=0.726; neuronal p<0.0001. Max caliper: astrocytic p=0.516; neuronal p=0.017.

We further characterized the morphology of nLBs and CAL by studying regularity parameters of MGS-stained granules in βIII-tubulin-positive/GFAP-negative areas (corresponding to nLBs) and GFAP-positive/βIII-tubulin-negative areas (corresponding to CAL). Malin^KO^+p62^KO^ nLBs showed changes in circularity, eccentricity and maximal caliper with respect malin^KO^ nLBs (Figure 4C, Supplementary Figure 2). No significant differences in CAL morphology were detected between malin^KO^+p62^KO^ and malin^KO^ brains (Figure 4D, Supplementary Figure 2). Taken together, these results indicate that p62 is necessary for the correct packing of nLBs.

### Malin^KO^+p62^KO^ mice present neuroinflammation

Astrogliosis, microgliosis and increased expression of genes related to neuroinflammation are characteristic traits of malin^KO^ brains (13, 17, 18, 38). Given the importance of p62 in inflammatory responses (39), we studied the impact of p62 deletion on these processes. GFAP and IBA1 immunostainings showed an increase in reactive astrocytes and microglia in the hippocampi of malin^KO^ mice (Figure 5A, B), as previously described (13, 17, 40). Malin^KO^+p62^KO^ mice showed similar GFAP and IBA1 stainings (Figure 5A, B), thereby indicating that the deletion of p62 alone did not significantly alter the number of reactive astrocytes or microglia. We also examined the presence of A1 astrocytes, a subset of reactive neurotoxic astrocytes whose cytoplasm accumulates the inflammatory component protein C3 (41). Malin^KO^+p62^KO^ mice showed a significant increase in reactive C3-positive astrocytes compared to control animals (Figure 5C, D), although they did not show a significant difference with malin^KO^ mice.

**Figure 5.**
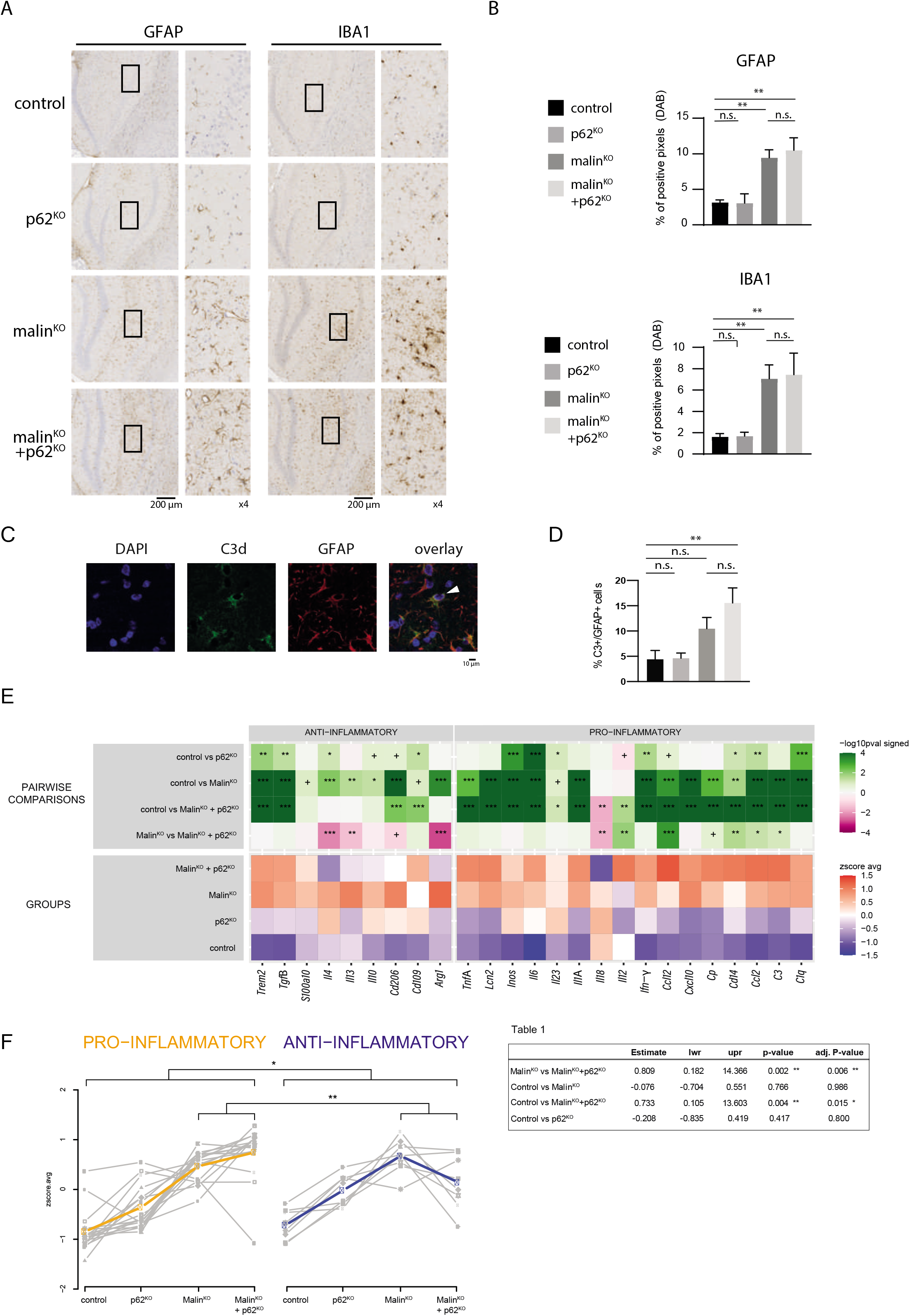
Neuroinflammation in malin^KO^ and malin^KO^+p62^KO^ mice. A-B. Representative immunostaining of GFAP (A) and IBA1 (B) in the hippocampus of the indicated experimental groups. C. Densitometry analysis of GFAP and IBA1 areas normalized to the hippocampal area. For comparisons between groups, one-way ANOVA followed by Holm’s Multiple Comparisons Test was performed using Prism7 software (GraphPad). Results are presented as the group mean ± SEM. n=5-6 animals per group. * P<0.05, ** P<0.01, *** P<0.001. *Adjusted p-values.* GFAP in hippocampus: control vs. p62^KO^ p= 0.9562; control vs. malin^KO^ p=0.0055; control vs. malin^KO^+p62^KO^ p<0.0028; and malin^KO^ vs. malin^KO^+p62^KO^ p=0.7931. GFAP in cortex: control vs. p62^KO^ p=0.5388; control vs. malin^KO^ p=0.5388; control vs. malin^KO^+p62^KO^ p=0.1823; and malin^KO^ vs. malin^KO^+p62^KO^ p=0.5388. IBA1 in hippocampus: control vs. p62^KO^ p=0.9719; control vs. malin^KO^ p=0.0175; control vs. malin^KO^+p62^KO^ p=0.0175; and malin^KO^ vs. malin^KO^+p62^KO^ p=0.9705. IBA1 in cortex: control vs. p62^KO^ p=0.9620; control vs. malin^KO^ p=0.8390; control vs. malin^KO^+p62^KO^ p=0. 8390; and malin^KO^ vs. malin^KO^+p62^KO^ p=0.9620. D. Representative super-resolution images of the immunofluorescence staining against C3d and GFAP in the hippocampus of a MalinKO mouse. One A1 reactive astrocyte is indicated in the arrohead. D. Quantification of the percentage of GFAP+ cells that are C3+ in the hippocampus. N=7-9 mice per group. * P<0.05, ** P<0.01, *** P<0.001. *Adjusted p-values.* control vs. p62^KO^ p>0.9999; control vs. malin^KO^ p=0.2539; control vs. malin^KO^+p62^KO^ p=0.0053; and malin^KO^ vs. malin^KO^+p62^KO^ p=0.3513. E. Heat map of pro- and anti-inflammatory transcripts quantified by qPCR normalized to the housekeeper gene GAPDH. N=6 animals per group. Linear mixed-effects models were fitted using ΔCt as response variable, the genotype as covariate of interest, the mouse experimental group replicate as adjusting factor and the mouse Id as random effect to account for variability in technical replicates. Comparisons were done independently for each gene measured. * P<0.05, ** P<0.01, *** P<0.001. Raw and adjusted p-values are summarized in Supplementary file 3. F. Left panel: dot plots of the average Z-score of the genes shown in panel A in the indicated experimental groups. Gray lines are indicative of one gene. Colored lines are the average of gray lines. Right panel (Table 1): linear mixed effect model that quantified consistency in ΔCts between genes of the same inflammatory group, across the 4 genotypes. * P<0.05, ** P<0.01, *** P<0.001.

We next examined the transcriptional profiles of cytokines and other mediators of the immune response associated with activated microglia. Malin^KO^ and malin^KO^+p62^KO^ mice showed a similar increase, with respect to control mice, in the expression of IL1-α, TNF-α and C1q, cytokines, key mediators of A1 astrocyte activation (41) (Figure 5E). We observed a similar result in other inflammation-associated genes, including LCN2, CXCL10, CCL12 and CCL2 (38, 42). The expression of genes involved in suppressing inflammation, like S100a10, IL10 and Arg-1A, was significantly increased in malin^KO^ mice (Figure 5E). Overall, malin^KO^ mice showed the upregulation of both anti- and pro-inflammation genes, while malin^KO^+p62^KO^ animals had a more marked increase of pro-inflammatory cytokines (IL12, CCL12, CD14, CCL2 and C3), and a reduced expression of the anti-inflammatory molecule Arg-1 and anti-inflammatory cytokines such as IL4 and IL13 (Figure 5E). Next, we compared normalized ΔCt patterns across the four genotypes for gene signatures defining pro- and anti-inflammatory activity. Interestingly, malin^KO^+p62^KO^ mice showed higher expression of the pro-inflammatory signature than malin^KO^ mice, whereas this trend was reversed for the anti-inflammatory signature, with malin^KO^ mice showing greater expression than malin^KO^+p62^KO^ counterparts. This pattern was consistent for most of the genes that defined the two signatures (Figure 5F). Overall, malin^KO^ and malin^KO^+p62^KO^ mice showed a similar inflammatory response, albeit modestly exacerbated in the latter.

### Deletion of p62 enhances the epileptic phenotype of malin^KO^ mice

Malin^KO^ mice present increased susceptibility to kainate-induced epilepsy (17, 30), a finding consistent with one of the main symptoms of LD patients. We have proposed that this pathological trait is due to the accumulation of LBs in neurons (13, 30). Therefore, we next aimed to examine whether the change in nLB morphology in malin^KO^+p62^KO^ brains was translated into a worsening of the epileptic phenotype. To that end, 5-month-old mice received three consecutive kainate injections (6 mg/kg, i.p. every 30 min) and were video-recorded for 240 minutes after the first injection to monitor behavior (i.e., epileptic events). Malin^KO^ and malin^KO^+p62^KO^ animals showed a significant decrease in the average onset time of the first seizure (Figure 6A) and an increase in the number of seizures compared to control mice (Figure 6B). Furthermore, malin^KO^+p62^KO^ animals showed the highest number of all kinds of seizures per animal, and the percentage of mice that reached the most severe stage (VI) was significantly higher in this group (Figure 6B). While mice from the four genotypes reached all severity stages, only in the malin^KO^ and malin^KO^+p62^KO^ groups all the mice reached stage IV and showed an increased proportion of mice reaching stages V and VI (Figure 6C). Regarding the time spent in each stage, again malin^KO^+p62^KO^ mice showed the highest time spent in severe stages (Figure 6D), and the lowest in mild stages (I-III, not shown). Malin^KO^ and malin^KO^+p62^KO^ animals spent more time in stage IV, the latter group being statistically higher than the former and the only one to show significantly greater time spent in stage V when compared to control animals (Figure 6D). In summary, the increased susceptibility to kainate-induced epilepsy of malin^KO^ mice was further increased in malin^KO^+p62^KO^ mice.

**Figure 6.**
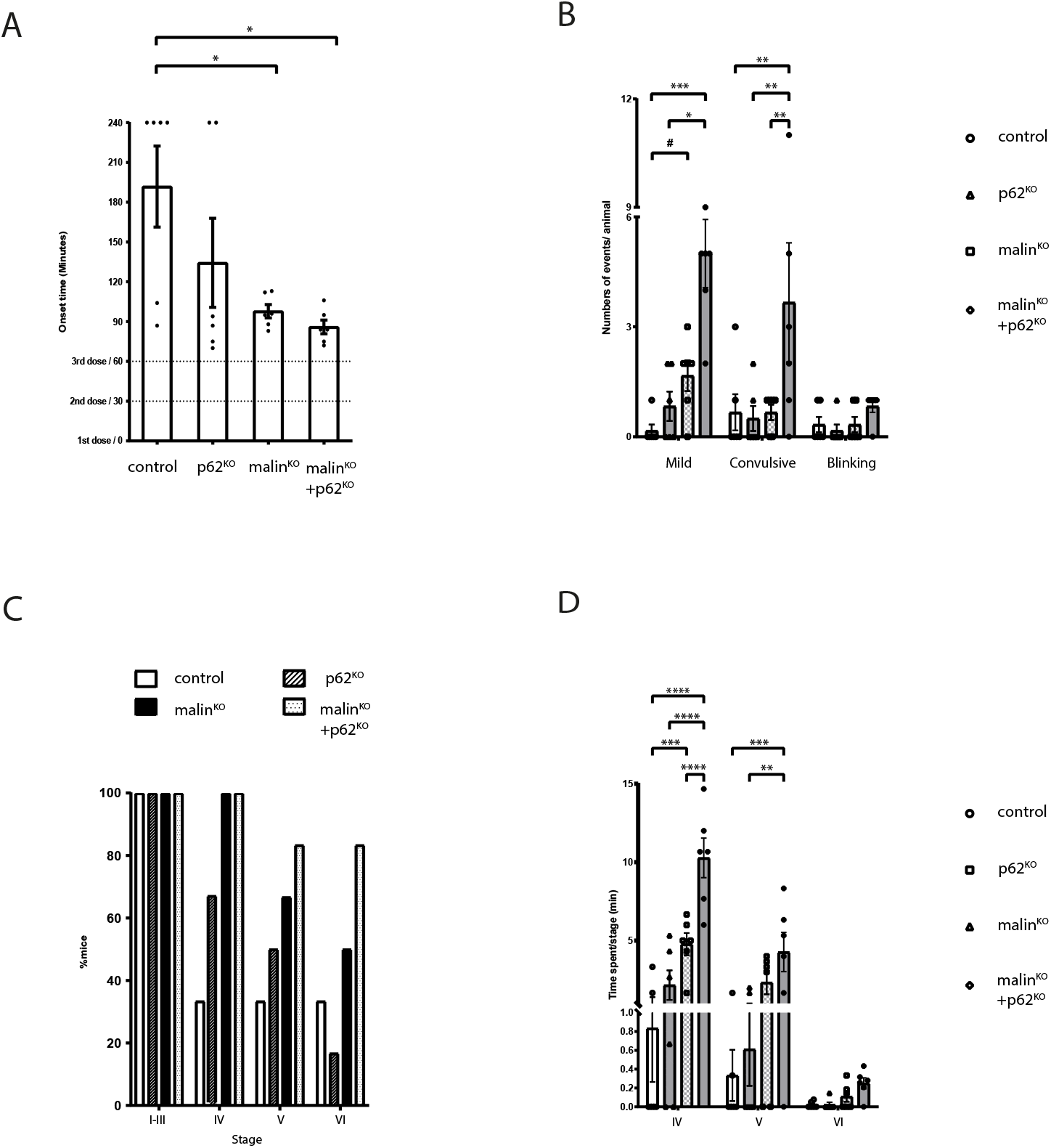
p62 deletion increases seizure susceptibility in malin^KO^ mice. A. Onset of epileptic activity in minutes. Data are expressed as average ±SEM. N=6 mice/group (*p<0.05) indicate significant difference (One-way ANOVA followed by Bonferroni test; p=0.0.0173). B. Number of seizures (mild, convulsive, or blinking) experienced per animal. Data are expressed as average ±SEM. N=6 mice/group. Two-way ANOVA (Seizure type factor: p=0.0003; Genotype factor: p=0.0012; interaction: p=0.0022). (*p<0.05, **p<0.01, ***p<0.005) indicate significant difference (Kruskal-Wallis for each seizure type followed by Dunn’s test); (#p<0.05) indicate pairwise differences between control and malin^KO^ by Mann-Whitney U-test. C. Percentage of mice reaching seizure stages I to VI. D. Time spent in each stage (expressed in minutes) during the course of the experiment. Data are expressed as average ±SEM. N= 6 mice/group. Two-way ANOVA (Stage factor (P <0.0001), the Genotype factor (P<0.0001), Interaction (P<0.0001)). (**p<0.01, ***p<0.005, ****p<0.001) indicate significant pairwise differences (post-hoc Bonferroni).

## DISCUSSION

The high levels of p62 in malin^KO^ brains (17, 18, 20) made us consider the possibility that its accumulation *per se* contributes to the etiopathology of LD. In this regard, it is worth noting that p62 interacts with many factors that play key roles in determining cell fate (26). The accumulation of p62 can lead to overactivation of oxidative stress responses through the Nrf2/Keap1 pathway (43) or can co-operate with other disease-associated proteins to induce cellular toxicity (44). Conversely, p62 depletion clears nuclear inclusion bodies and increases lifespan in a model of Huntington’s disease (29). Strikingly, our results indicate that the accumulation of p62 *per se* does not underlie the etiopathology of LD, as malin^KO^+p62^KO^ mice are not rescued from the characteristic pathological traits of LD but instead present exacerbated pathology.

p62 participates in the autophagic removal of protein aggregates in a process known as aggrephagy (45). p62 contains multiple protein-interaction domains, including a ubiquitin-binding domain and an oligomerization domain, which allow it to bind to and aggregate polyubiquitinated proteins into less harmful inclusions (46). The presence of p62 in LBs led us to hypothesize that glycogen aggregation in LD follows a similar pattern. Importantly, LBs were absent in the skeletal muscles and hearts of malin^KO^+p62^KO^ mice, thereby revealing that the abnormal glycogen that is formed in LD does not aggregate by itself, as it has been generally assumed, since p62 is essential for the formation of LBs in these tissues. These results have important implications for other diseases in which glycogen aggregates accumulate in skeletal muscle and heart, including Andersen’s disease (OMIM 232500) (47), Cori’s disease (OMIM 232400) (48), Tarui disease (OMIM 232800) (49), polyglucosan body myopathy-1 (OMIM 615895) (50) and polyglucosan body myopathy-2 (OMIM 616199) (51). In contrast, the brains of these animals still contained LBs, indicating that the process of LB formation is tissue-specific. In this regard, it is worth noting that aggrephagy genes appear to be differentially used in a tissue-specific manner (45). Functional redundancy of autophagy receptors (52, 53) could explain the presence of LBs in malin^KO^+p62^KO^ brains, as other receptors might compensate for the absence of p62 to trigger glycogen aggregation in this tissue.

However, super-resolution analysis revealed that p62 deletion results in more irregular, less round, less compact aggregates in neurons, thereby confirming the involvement of p62 in LB formation also in this cell type. In this regard, it is interesting to note that protein aggregates rich in p62 *(p62 bodies)* have liquid-like properties (high sphericity) and can undergo fusion events (54). Therefore, it is conceivable that the deletion of p62 blocks the liquid-like properties (roundness) and the fusion of insoluble glycogen aggregates into larger droplets. We have proposed that the epileptic phenotype of LD is due to the accumulation of glycogen in neurons (13, 30). Our results show that the change in the morphology of nLBs in malin^KO^+p62^KO^ mice is accompanied by an increase in susceptibility to kainate-induced epilepsy, thereby corroborating that the proper sequestration of abnormal glycogen into nLBs is essential to minimize its toxic effects in neurons.

Although p62 is present both in CAL and nLBs in the brains of malin^KO^ mice (12), analysis of the number and morphology of astrocytic CAL did not show significant differences in malin^KO^+p62^KO^ mice. However, the morphology of CAL is inherently heterogeneous (12), which could hamper the detection of changes in the parameters studied. Thus, we cannot discard that the deletion of p62 also affected CAL formation. We have recently demonstrated that the accumulation of CAL in astrocytes underlies neuroinflammation in LD (13). Accordingly, the cytokine inflammatory program was sustained and even switched toward a potential exacerbation of the neuroinflammation in malin^KO^+p62^KO^ mice. These results confirm the key role of astrocyte-driven inflammation in the pathophysiology of LD.

A longstanding question in LD is whether LBs are the toxic species themselves or whether they are formed to minimize the toxic consequences of the accumulation of abnormal glycogen, by sequestering it into less harmful aggregates. In this regard, there is wide consensus that in proteinopathies, early stages of aggregation are responsible for cellular toxicity and neurodegeneration (55–57). Malin^KO^+p62^KO^ mice, in which the formation of brain LBs is altered, offer the opportunity to study intermediate states of LB formation. We observed increased susceptibility to kainate-induced epilepsy in malin^KO^+p62^KO^ mice, which supports the hypothesis that immature glycogen aggregates are more toxic than mature LBs. On the basis of all these considerations and our findings, we propose a scenario in which LBs play a similar role as neurodegeneration-associated protein inclusion bodies (56, 58, 59). Poorly branched glycogen, which cannot be degraded by glycogen phosphorylase, would be formed as a side-product of glycogen metabolism. The malin/laforin complex would serve to prevent the formation of this abnormal glycogen. Thus, in the absence of laforin or malin, it would accumulate. In this context, p62 (together with other autophagy adaptors in the case of the brain) would promote its aggregation into LBs in order to minimize the toxic consequences of glycogen accumulation in neurons (19) and possibly in astrocytes (13) (Figure 7).

**Figure 7.**
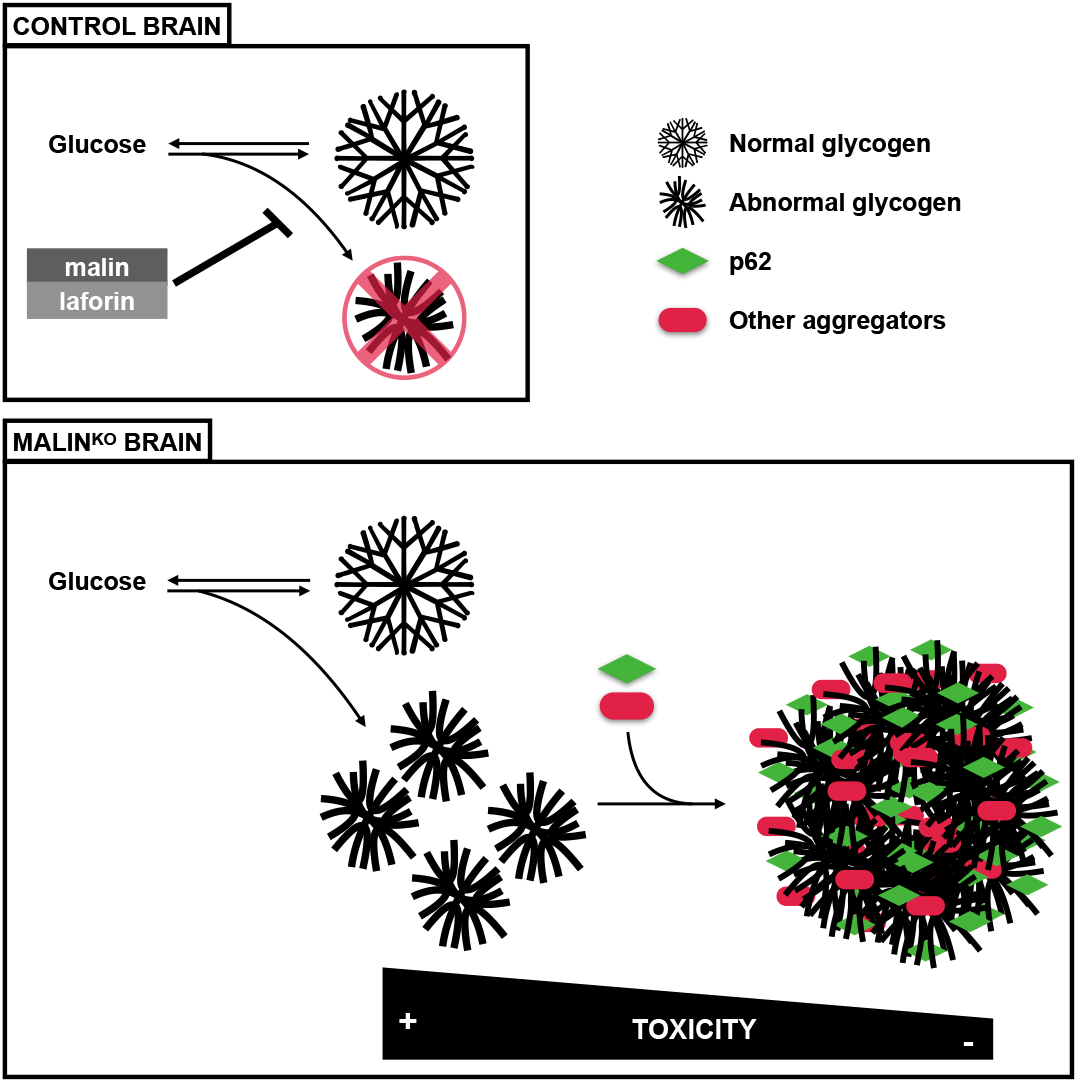
Graphical summary of the study. Abnormal glycogen is produced as a side-product of glycogen metabolism. The malin/laforin complex prevents the formation of this abnormal polysaccharide. In the absence of laforin or malin, abnormal glycogen accumulates. In this context, p62 promotes its aggregation into LBs, to minimize the toxic consequences of its accumulation. p62 deletion correlates with an exacerbation of the inflammation and epilepsy supporting the notion that LBs are neuroprotective.

Beyond LD, we have witnessed an increasing body of literature showing glycogen accumulation as a new common thread in aging, neurodegenerative diseases (including Alzheimer’s, Parkinson’s and Huntington’s disease and amyotrophic lateral sclerosis (60–62)) and epilepsy (63–66). Moreover, the accumulation of *corpora amylacea,* p62 and aggregation-prone proteins in the brain correlates with aging and neurodegeneration (8, 14, 67, 68). The accumulation of glycogen in these conditions could play an active role in disrupting cell homeostasis, causing neuroinflammation and epilepsy, as we have shown for LD. Therefore, glycogen synthesis emerges as a potential therapeutic target for aging and other neurological diseases.

## Conclusions

Our study shows that p62 is necessary for LB formation in muscle and heart and that it is also important for the correct formation of LBs in the brain. The deletion of p62 worsens the pathology of LD. Our results provide an unprecedented description of a protective role of p62-directed sequestration of glycogen aggregates in LD.

## Supporting information

Sumpplemental files

## LIST OF ABBREVIATIONS

Arg-1: arginase-1
C1q: complement component 1q
C3: complement component 3
CA: *corpora amylacea*
CAL: *corpora amylacea*-like bodies
CCL2: chemokine (C-C motif) ligand 2
CD14: cluster of differentiation 14
CXCL10: C-X-C motif chemokine ligand 10
CCL12: chemokine ligand 12
EPM2a: Epilepsy, Progressive Myoclonus Type 2A
EPM2b: Epilepsy, Progressive Myoclonus Type 2b
GFAP: glial fibrillary acidic protein
IBA1: ionized calcium-binding adapter molecule 1
IL1-α: interleukin 1 α
IL10: interleukin 10
IL12: interleukin 12
IL13: interleukin 13
IL4: interleukin 4
KA: Kainic acid
LBs: Lafora bodies
LCN2: lipocalin-2
LD: Lafora disease
MGS: muscle glycogen synthase
NHLRC1: _NHL Repeat Containing E3 Ubiquitin Protein Ligase 1
nLBs: neuronal Lafora bodies
PAS: periodic acid-Schiff
S100A10: S100 calcium-binding protein A10
TNF-α: tumor necrosis factor α

## DECLARATIONS

### Ethics approval and consent to participate

All procedures were approved by the Barcelona Science Park’s Animal Experimentation Committee and were carried out in accordance with the European Community Council Directive and National Institutes of Health guidelines for the care and use of laboratory animals.

### Consent for publication

Not applicable.

### Availability of data and materials

The data that support the findings of this study are available from the corresponding author upon reasonable request.

## AUTHORS’ CONTRIBUTIONS

PP and JD designed and performed the main experiments and analyzed the data. AH and JADR conducted the in-vivo experiments. OV, ILS, MKB, AG, MA and NP conducted additional experiments. PP wrote the manuscript. JD and JJG designed the project and edited the manuscript. All authors have read and approved the final version of the manuscript.

## ACKNOWLEDGMENTS

We thank Anna Adrover and Vanessa Fernandez for technical assistance and IRB Barcelona’s Advanced Digital Microscopy Facility, Histopathology Facility and Bioinformatics and Biostatistics Facility for technical support. Thanks also go to Nikos Giakoumakis for his kind support with super-resolution microscopy and Adrià Caballé for support with statistical analysis. Tanya Yates for correcting the English manuscript. p62^KO^ mice were kindly provided by Dr. Maria Sonegas (Spanish National Cancer Research Centre-CNIO, Madrid, Spain) with permission from Prof. Tetsuro Ishii (University of Tsukuba, Japan).

## SUPPLEMENTARY FIGURES

**Supplementary Figure 1. p62 deletion prevents glycogen aggregation in heart.**

A. Histological localization of LBs in heart. Periodic acid-Schiff staining (PAS) is shown for the indicated tissues of 11-month-old malin^KO^ and 11-month-old malin^KO^+p62^KO^ littermates. Scale bar=25 μ m. B. Representative immunostaining of p62 and MGS in heart. Scale bar=25 μm

**Supplementary Figure 2. Additional morphology parameters of neuronal LBs and astrocytic CAL.**

Quantifications of the indicated morphological parameters in neuronal LBs and astrocytic CAL. A linear mixed model analysis was performed for statistical analysis. N=6 animals per genotype. *Adjusted p-values.* Area: astrocytic p=0.657; neuronal p=0.269. Perimeter: astrocytic p=0.952; neuronal p=0.592. MGS mean intensity: astrocytic p=0.771; neuronal p=0.388. Centroid Xm: astrocytic p=0.392; neuronal p=0.959. Centroid Ym: astrocytic p=0.646; neuronal p=0.809. Min caliper: astrocytic p=1; neuronal p=0.162.

**Supplementary File 3.**

Raw and adjusted p-values of the pairwise comparisons in Figure 7A.

**Supplementary File 4.**

SYBR green primers used in Figure 7.

