## Supplementary material for "Lack of p62 impairs glycogen aggregation and exacerbates pathology in a mouse model of myoclonic epilepsy of Lafora": Sumpplemental files

### Supplementary Figure 1

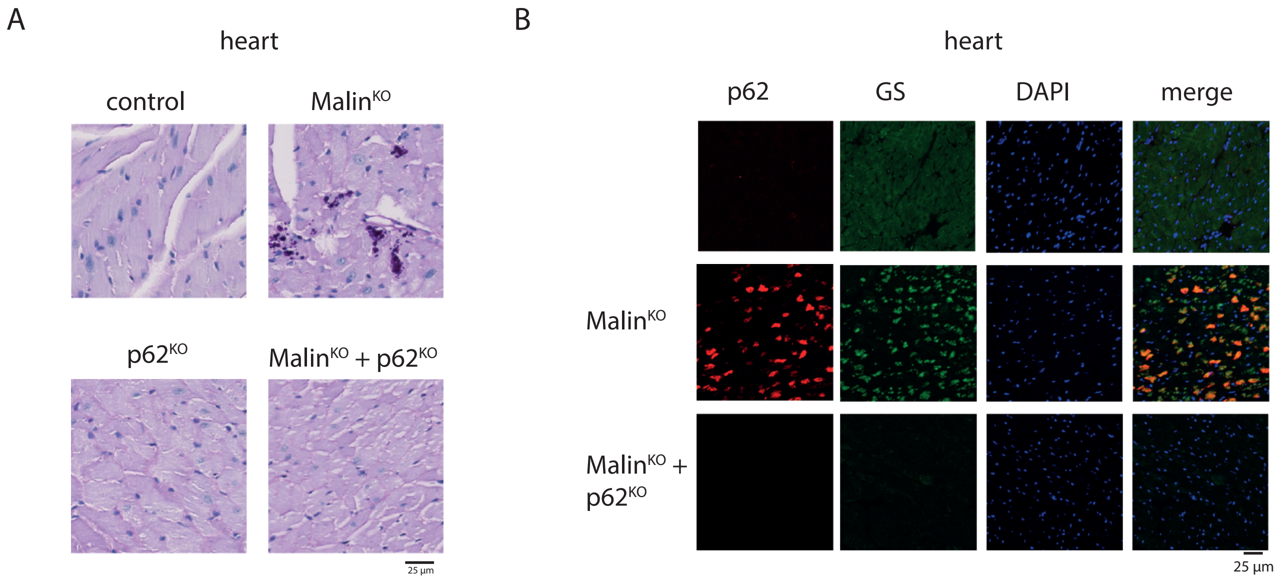

### Supplementary Figure 2

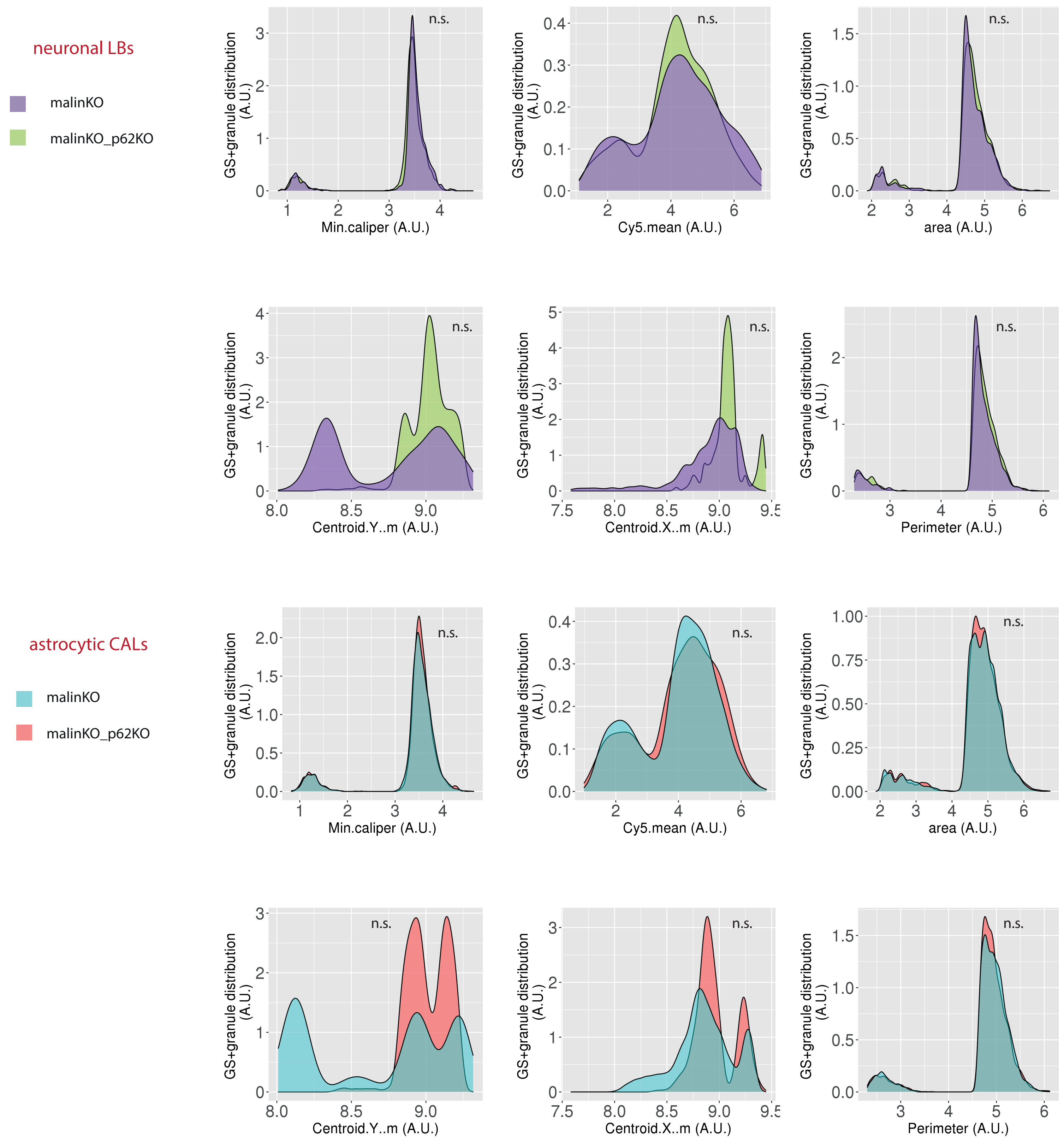

### Supplementary File 3

**raw p-values**

| gene | p62KO -CTRL | malinKO -CTRL | malinKO+p62KO -CTRL | malinKO+p62KO -malinKO |
|---|---|---|---|---|
| iNOS | 0,000627985 | 3,28E-05 | 3,72E-05 | 0,81796078 |
| IL6 | 3,83E-06 | 7,50E-09 | 7,02E-13 | 0,29406075 |
| IFN-g | 0,024965437 | 8,09E-05 | 1,49E-06 | 0,576956352 |
| C1q | 0,002368484 | 1,66E-07 | 8,23E-09 | 0,804516837 |
| TREM2 | 0,013795538 | 9,47E-10 | 1,56E-10 | 0,975476864 |
| IL12 | 0,180660635 | 0,700943651 | 0,025707614 | 0,012892388 |
| IL18 | 0,649799372 | 0,925090468 | 0,019875824 | 0,020941161 |
| IL23 | 0,096005457 | 0,150064159 | 0,081135417 | 0,826105466 |
| CD206 | 0,183028442 | 2,12E-05 | 0,005927228 | 0,101948059 |
| IL4 | 0,078967515 | 0,00756228 | 0,930594325 | 0,005889722 |
| IL10 | 0,172617973 | 0,051169739 | 0,304207368 | 0,457743055 |
| TGFb | 0,045404599 | 1,98E-05 | 3,61E-07 | 0,566873938 |
| C3 | 0,257200748 | 4,25E-13 | 0 | 0,096674678 |
| CCL2 | 0,04324087 | 8,49E-11 | 0 | 0,061833936 |
| CXCL10 | 0,855087833 | 1,21E-10 | 8,90E-10 | 0,75650904 |
| CXCL12 | 0,182336696 | 0,000693553 | 1,25E-11 | 0,000732561 |
| LCN2 | 0,419700063 | 2,49E-08 | 3,82E-10 | 0,492117664 |
| Arg1 | 0,868773282 | 0,00041276 | 0,751981875 | 0,001300992 |
| Cd109 | 0,050671929 | 0,123805406 | 0,008760502 | 0,279134297 |
| Cd14 | 0,076345881 | 0,020858112 | 7,18E-06 | 0,029430864 |
| Cp | 0,987136866 | 0,002233728 | 1,56E-05 | 0,18413109 |
| S100a10 | 0,785335781 | 0,13050126 | 0,277190034 | 0,670490581 |
| TNFa | 0,719892712 | 0,002766171 | 6,65E-05 | 0,319335641 |
| IL13 | 0,687529219 | 0,034313818 | 0,957773761 | 0,038828346 |
| IL1b | 3,05E+05 | 7,94E+00 | 6,76E-01 | 6,16E+05 |

**adjusted p-values**

| gene | p62KO -CTRL | malinKO -CTRL | malinKO+p62KO -CTRL | malinKO+p62KO -malinKO |
|---|---|---|---|---|
| iNOS | 0,00224542 | 0,00012611 | 0,00017007 | 0,99367684 |
| IL6 | 1,10E-05 | 2,38E-08 | 1,48E-12 | 0,65056 |
| IFN-g | 0,0833311 | 0,0003818 | 3,79E-06 | 0,922294 |
| C1q | 0,0089913 | 4,52E-07 | 2,41E-08 | 0,9923234 |
| TREM2 | 0,047681 | 2,01E-09 | 4,17E-10 | 0,999984 |
| IL12 | 0,460676 | 0,972368 | 0,085889 | 0,0447 |
| IL18 | 0,955528 | 0,999558 | 0,067333 | 0,070732 |
| IL23 | 0,27596 | 0,39789 | 0,23919 | 0,99449 |
| CD206 | 0,469956 | 7,20E-05 | 0,021667 | 0,294061 |
| IL4 | 0,23363 | 0,027158 | 0,999648 | 0,021412 |

| | | | | |
|---|---|---|---|---|
| IL10 | 0,44436 | 0,15988 | 0,66558 | 0,83882 |
| TGFb | 0,14353 | 7,65E-05 | 2,76E-06 | 0,91657 |
| C3 | 0,59361 | 1,10E-12 | < 2.22e-16 | 0,27707 |
| CCL2 | 0,13716 | 1,95E-10 | < 2.22e-16 | 0,18856 |
| CXCL10 | 0,99678 | 2,68E-10 | 2,05E-09 | 0,98483 |
| CXCL12 | 0,4620595 | 0,002663 | 2,76E-11 | 0,0027137 |
| LCN2 | 0,80204 | 9,52E-08 | 5,40E-09 | 0,86601 |
| Arg1 | 0,997609 | 0,001512 | 0,983967 | 0,004792 |
| Cd109 | 0,158076 | 0,340153 | 0,031052 | 0,627567 |
| Cd14 | 0,22646 | 0,070539 | 2,22E-05 | 0,096618 |
| Cp | 0,9999977 | 0,0082964 | 5,31E-05 | 0,4609317 |
| S100a10 | 0,98958 | 0,3551 | 0,62464 | 0,96266 |
| TNFa | 0,97696141 | 0,00996043 | 0,00024165 | 0,68483843 |
| IL13 | 0,9683 | 0,11141 | 0,99992 | 0,12496 |
| IL1b | 0.66467 | 3,96E-01 | 2,64E-02 | 0.94127 |

### Supplementary File 4

| TARGET | FORWARD | REVERSE |
|---|---|---|
| TREM2 | GACCTCTCCACCAGTTTCTCC | TACATGACACCCTCAAGGACTG |
| TGFb | TGACGTCACTGGAGTTGTACGG | GGTTCATGTCATGGATGGTGC |
| S100a10 | CTGCTCACAAGAAGCAGTGG | CCTCTGGCTGTGGACAAAAT |
| IL-4 | GGTCTCAACCCCCAGCTAG | GCCGATGATCTCTCTCAAGTGAT |
| IL13 | GGGTGACTGCAGTCCTGGCT | GTTGCTCAGCTCCTCAATAAGC |
| IL10 | CCAAGCCTTATCGGAAATGA | TTTTCACAGGGGAGAAATCG |
| CD206 | CAAGGAAGGTTGGCATTTGT | CCTTTCAGTCCTTTGCAAGC |
| Cd109 | GCAGCGATTTCGATGTCCAC | CACAGTCGGGAGCCCTAAAG |
| Arg1 | CCTCGAGGCTGTCCTTTTGA | TTTTAGGGTTACGGCCGGTG |
| Tnfa | CTTGATGGTGGTGCATGAGA | TGTGCTCAGAGCTTTCAACAA |
| Lcn2 | CACACTCACCACCCATTCAG | CCAGTTCGCCATGGTATTTT |
| iNOS | GGCAGCCTGTGAGACCTTTG | CATTGGAAGTGAAGCGTTTCG |
| IL6 | TCCTACCCCAACTTCCAATGCTC | TTGGATGGTCTTGGTCCTTAGCC |
| IL23 | TGTGCCCCGTATCCAGTGT | CGGATCCTTTGCAAGCAGAA |
| IL1β | ACGGACCCCAAAAGATGAAG | TTCTCCACAGCCACAATGAG |
| IL12 | GGAAGCACGGCAGCAGAATA | AACTTGAGGGAGAAGTAGGAATGG |
| IFN-γ | TCAAGTGGCATAGATGTGGAAGAA | TGGCTCTGCAGGATTTTCATG |
| CXCL10 | GACCATCAAGAATTTAATGAAAGCG | CCATCCACTGGGTAAAGGG |
| Cp | AGTGTATAGAGGATGTTCCAGGTCA | TGTGATGGGAATGGGCAATGA |
| Cd14 | GCTTCAGCCCAGTGAAAGAC | GGACTGATCTCAGCCCTCTG |
| CCL2 | GAGTAGGCTGGAGAGCTACAAGAG | AGGTAGTGGATGCATTAGCTTCAG |
| C1q | CCCTGGTAAATGTGACCCTTTT | TCTGCACTGTACCCGGCTA |
| C3 | TCCTGAACTGGTCAACATGG | AAACTGGGCAGCACGTATTC |
| Ccl12 | ACCATCAGTCCTCAGGTATTGG | TTCCGGACGTGAATCTTCTG |
| IL18 | GCCTCAAACCTTCCAAATCA | TGGATCCATTTCCTCAAAGG |
